# Cross-species optimization of nuclei isolation in plants

**DOI:** 10.1101/2025.09.04.674249

**Authors:** Yun Luo, Jiali Yan, Thuy La, Jianbing Yan, M. Cinta Romay

**Affiliations:** Institute for Genomic Diversity, Cornell University, Ithaca, NY 14853, USA; National Key Laboratory of Crop Genetic Improvement, Huazhong Agricultural University, Wuhan 430070, China; Hubei Hongshan Laboratory, Huazhong Agricultural University, Wuhan 430070, China

## Abstract

Single-cell technologies are transforming plant biology, yet broadly transferable nuclei isolation remains a key bottleneck for snRNA-seq. We developed a reproducible, Percoll-based workflow that is applicable to multiple maize tissues and eight additional plant species. In maize, nuclei from root, stem, leaf, and embryo consistently concentrated at the 80% Percoll interface and exhibited high integrity, with typical recoveries > 50,000 nuclei per sample. For other species, gradient compositions were tuned according to genome size to achieve efficient enrichment and clean suspensions, and yields ranged from 20,000 to 50,000 nuclei per sample. Downstream validation showed that nuclei from special interest maize and *Tripsacum* generated high-quality 10x Genomics snRNA-seq libraries, as supported by cDNA quality profiles. These results demonstrate the versatility and robustness of the method across species and tissues.

## Introduction

Plants are highly complex organisms composed of a wide variety of specialized cell types that coordinate distinct biological functions in a spatial and temporal manner [1–5]. Traditional bulk RNA sequencing (RNA-seq) approaches typically analyze entire tissues, organs, or even whole plants [6], thereby averaging gene expression across multiple cell types and masking cell-specific transcriptional programs [7]. To overcome these limitations, single-cell and single-nucleus RNA sequencing (sc/snRNA-seq) technologies have emerged as powerful tools for dissecting cellular heterogeneity and enabling high resolution, cell-type-specific transcriptomic profiling in plants [3,5,8–13]. In plants, sc/snRNA-seq typically involves two main strategies: enzymatic digestion of cell walls to isolate protoplasts for scRNA-seq, and detergent-based lysis of the plasma membrane to extract nuclei for snRNA-seq [10,14,15]. Compared with protoplast-based scRNA-seq, nuclei-based methods offer key advantages: they minimize bias against hard-to-dissociate or fragile cell types, avoid stress-induced transcriptional artifacts caused by enzymatic digestion, and are compatible with frozen, lignified, or otherwise recalcitrant tissues [10,16–18].

Despite these advantages, isolating intact and high-purity nuclei from plant tissues remains technically challenging [19–21]. The presence of rigid cell walls, abundant secondary metabolites, and high tissue heterogeneity complicates buffer penetration, nuclear release, and debris removal, often resulting in low yields or damaged nuclei [10,22–24]. To address these issues, density gradient centrifugation is widely employed to enrich intact nuclei from crude lysates based on buoyant density [5,25–29]. Among available media, Percoll - a colloidal silica solution coated with polyvinylpyrrolidone (PVP) - is especially well-suited for plant nuclei isolation because it forms iso-osmotic, low-viscosity gradients that allow gentle and efficient separation of nuclei from cytoplasmic debris, starch granules, and organelles [30,31]. Furthermore, Percoll gradients have been shown to reduce ribonuclease activity associated with nuclear fractions, thereby improving RNA preservation. During centrifugation, nuclei migrate to specific layers based on size and density, typically forming a distinct milky-white band that facilitates visual identification and collection. Importantly, nuclear buoyancy is influenced by genome size, making it necessary to empirically adjust Percoll gradient compositions for different species and tissue types [32]. Despite its broad use, there is still a lack of standardized and comparative protocols for nuclei isolation across species with different genome sizes and tissue characteristics.

In this study, we developed a robust, broadly applicable nuclei isolation protocol optimized for a range of plant species and tissues. We tested this protocol across nine species - including both monocots and dicots - with genome sizes ranging from 0.135 to 16 Gb. By tailoring Percoll gradient compositions to match genome size and tissue characteristics, we successfully extracted intact nuclei from roots, leaves, stems, and embryos. Nuclear integrity and purity were confirmed using DAPI staining, microscopy, and automated nuclei counting. Furthermore, we validated the suitability of our preparations for downstream applications by generating high-quality cDNA libraries from maize and *Tripsacum* leaf nuclei using the 10x Genomics Chromium platform. Our results demonstrate the versatility and effectiveness of this method for plant single-nucleus transcriptomics, and provide practical guidance for nuclei isolation across diverse plant systems.

## Methods

### Plant materials

In this study, we investigated nine species: maize (*Zea mays*. L), Tripsacum (*Tripsacum dactyloides*), barely (*Hordeum vulgare*), oat (*Avena sativa*), winter wheat (*Triticum aestivum*), rice (*Oryza sativa*), sorghum (*Sorghum bicolor*), Arabidopsis (*Arabidopsis thaliana*), and Blue grama (*Bouteloua gracilis*). All plants were grown in the Cornell University greenhouse under controlled conditions with a 14-hour photoperiod of supplemental light and a day/night temperature regime of 28 °C/22 °C. For leaf samples, 0.5 g fresh leaf tissue was collected for nuclei isolation 10-14 days after germination in maize, oat, winter wheat, rice, and sorghum and 25-30 days after sowing in Arabidopsis, blue grama, and *Tripsacum*. For root samples, maize, *Tripsacum*, barely, oat, winter wheat, rice, sorghum, and blue grama were germinated on moist germination paper in the greenhouse, and 1-2 cm root tips were collected for nuclei isolation when roots reached ∼ 5 cm in length (4-10 days). Arabidopsis were grown vertically on ½ MS agar medium for 7-10 days, and root tips (∼5 cm) were similarly collected for nuclei isolation. In addition, maize stem tissue was collected from the same seeding at the time of root sample, and maize embryos were harvested 15 days after pollination for nuclei isolation.

### Chemical and stock solutions

1. 5 M KOH: Dissolve 14.05 g potassium hydroxide (KOH, Sigma-Aldrich, #221473-500G) in 40 mL RNase-free water and brought up to 50 mL. The solution was sterile-filtered (0.22μm) and stored at 4°C.
2. 5 M NaOH: A total of 10 g sodium hydroxide (NaOH, Sigma-Aldrich, #221465-500G) was fully dissolved in 40 mL RNase-free water. The final volume was adjusted to 50 mL, followed by filtration (0.22 μm) and storage at 4 °C.
3. 2 M HCl: 8.32 mL concentrated Hydrochloric acid (HCl, Sigma-Aldrich, #258148-500ML) was carefully added to 30 mL of RNase-free water under a fume hood. After cooling to room temperature, the mixture was topped up to 50 mL and sealed for storage at 4 °C.
4. 1.25 M MES (pH 5.6-6.7):12.2 g 2-(N-morpholino) ethanesulfonic acid (MES, VWR chemicals, #E183-100G) power was mixed with 30 mL RNase-free water. The PH was adjusted using 5 M KOH, then brought volume to 50 ml. The buffer was passed through a 0.22 μm filter and stored at 4 °C. (see Note 1).
5. 3 M KCl: 11.18 g potassium chloride (KCl, Sigma-Aldrich, #P9541-500G) was dissolved in 40 mL RNase-free water and adjusted to 50 mL. After filtration (0.22 μm), it was kept at 4 °C.
6. 1 M MgCl_2_: Dissolve 4.76 g anhydrous magnesium chloride (MgCl_2_, Invitrogen, #AM9530G) in 40 ml RNase-free water, and made up to 50 mL. The solution was sterilized by filtration (0.22 μm) and stored at 4 °C.
7. 20% Triton X-100: 2 mL Triton X-100 (Sigma-Aldrich, #T8787-100ML) was combined with 8 mL RNase-free water to yield a 20% (v/v) working solution. The mixture was stored at 4 °C.
8. 1 M DTT: 1.5425 g Dithiothreitol (DTT; Roche, #10708984001-10G) was dissolved in 10 mL RNase-free water, sterile-filtered, aliquoted (1 mL), and frozen at –20 °C in light-protected tubes (see Note 2).
9. 1× PBS (pH = 7.4): To prepare 10× PBS (50 mL), dissolve 4 g sodium chloride (Nacl, Sigma-Aldrich, #S5886-500G), 0.1 g KCl, 0.72 g sodium phosphate dibasic (Na_2_HPO_4_, Millipore, #567547-250G) and 0.12 g potassium phosphate monobasic (KH_2_PO_4_, Millipore, #529586-250G) in 40 mL RNase-free water. Adjust the pH to 7.4 using 1 M NaOH or HCl, then bring to 50 mL and filter (0.22 μm). Then, to obtain 1× PBS, dilute 5 mL of 10× stock with 45 mL RNase-free water. Store at 4 °C.
10. 100 mM Spermine tetrahydrochloride: A total of 69.65 mg spermine tetrahydrochloride (Sigma-Aldrich, #S1141-5G) was dissolved in RNase-free water to a final volume of 2 mL. Aliquots were stored at –20 °C (see Note 2).
11. 100 mM Spermidine trihydrochloride: 50.9 mg spermidine trihydrochloride (Sigma-Aldrich, S2501-5G) was dissolved in RNase-free water and adjusted to 2 mL. Store at –20 °C (see Note 2).
12. 5 ng/μL DAPI Solution: Dilute 5 μL of 1 mg/mL 4′,6-diamidino-2-phenylindole (DAPI, Roche, #10236276001-10MG) stock with 995 μL 1 × PBS. Mix thoroughly and store at –20 °C in the dark (see Note 2).
13. 100 mM PMSF: 17.42 mg Phenylmethylsulfonyl fluoride (PMSF, Roche, #10837091001) was freshly dissolved in 1 mL absolute ethanol. The solution was used within one week and stored at –20 °C (see Note 3).
14. 1% BSA: 0.1 g bovine serum albumin (BSA, VWR Life Science, #0903-5G) was gradually added to 8 mL RNase-free water with gentle stirring. After complete dissolution, the volume was brought to 10 mL, mixed thoroughly, aliquoted, and stored at –20 °C.
15. Percoll (MP Biomedicals, #MF0219536990): Stored at 4 °C and gently mix before use (see Note 4).
16. 2-Methyl-2,4-pentanediol (Sigma-Aldrich, #8.20819-1L): Stored at temperature.
17. RNase-free water (Invitrogen, #10977015): Stored at 4 °C and used in aliquots.
18. Plant Protease Inhibitors (Thermo Scientific Chemicals, #J64576.XF): Stored at –20 °C.
19. Protector RNase Inhibitor 40U/ μL (Roche, #41181611): Stored at –20 °C.

### Solutions and Media

Prepare pre-extraction stock solutions and daily-use working solutions as shown in Table 1 and Table 2, respectively (see Note 5).

**Table 1.** Pre-extraction stock solutions for nuclei isolation and purification. Refer to Notes 1 to 5 for storage condition.

| # | Buffer Components | Stock solution | Volume |
| --- | --- | --- | --- |
| **A** | **4 x EB (total 9 mL)** |  |  |
| 1 | 24%(v/v) 2-Methyl-2,4-pentanediol | 2-Methyl-2,4-pentanediol | 2.16 mL |
| 2 | 80 mM MES-KOH (pH 5.7-6.7) | 1.25 M MES-KOH (pH 5.7-6.7) | 576 µL |
| 3 | 80 mM KCl | 3 M KCl | 240 µL |
| 4 | 40 mM MgCl<sub>2</sub> | 1 M MgCl<sub>2</sub> | 360 µL |
| 5 | RNase-free water | RNase-free water | 5.664 mL |
| **B** | **1 x EBM (total 12 mL)** |  |  |
| 1 | 0.25V total | 4 × EB | 3 mL |
| 2 | RNase-free water | RNase-free water | 9 mL |
| **C** | **10 x GRA (total 50 mL)** |  |  |
| 1 | 60%(v/v) 2-Methyl-2,4-pentanediol | 2-Methyl-2,4-pentanediol | 30 mL |
| 2 | 0.2 M MES-KOH (pH 5.7-6.7) | 1.25 M MES-KOH (pH 5.7-6.7) | 8.48 mL |
| 3 | 0.2 M KCl | 3 M KCl | 3.333 mL |
| 4 | 0.1 M MgCl<sub>2</sub> | 1 M MgCl<sub>2</sub> | 5 mL |
| 5 | RNase-free water | RNase-free water | 3.187 mL |
| **D** | **1× GRAM (total 50 mL)** |  |  |
| 1 | 0.1V total | 10 x GRA | 5mL |
| 2 | RNase-free water | RNase-free water | 45 mL |
| **E** | **100% PM (total 50 mL)** |  |  |
| 1 | 0.9V total | Percoll | 45 mL |
| 2 | 0.1V total | 10 x GRA | 5 mL |
| **F** | **90% PM (total 20 mL)** |  |  |
| 1 | 0.9V total | 100% PM | 18 mL |
| 2 | 0.1V total | 1× GRAM | 2 mL |
| **G** | **80% PM (total 20 mL)** |  |  |
| 1 | 0.8V total | 100% PM | 16 mL |
| 2 | 0.2V total | 1× GRAM | 4 mL |
| **H** | **75% PM (total 20 mL)** |  |  |
| 1 | 0.75V total | 100% PM | 15 mL |
| 2 | 0.25V total | 1× GRAM | 5 mL |
| **I** | **70% PM (total 20 mL)** |  |  |
| 1 | 0.7V total | 100% PM | 14 mL |
| 2 | 0.3V total | 1× GRAM | 6 mL |
| **J** | **50% PM (total 20 mL)** |  |  |
| 1 | 0.5V total | 100% PM | 10 mL |
| 2 | 0.5V total | 1× GRAM | 10 mL |
| **K** | **25% PM (total 20 mL)** |  |  |
| 1 | 0.25V total | 100% PM | 5 mL |
| 2 | 0.75V total | 1× GRAM | 15 mL |

**Table 2.** Daily-use working solutions for nuclei isolation and purification. See **Notes 5, 6** for detailed instructions.

| # | Buffer Components | Stock solution | Volume |
| --- | --- | --- | --- |
| **A** | **1 x EB\* (total 8 mL)** |  |  |
| 1 | 1× EBM | 1 x EBM | 8 mL |
| 2 | 1 mM DTT | 1 M DTT | 8 µL |
| 3 | 0.2% TritonX-100 | 20% TritonX-100 | 80 µL |
| 4 | RNase inhibitor 0.4U / µL | RNase inhibitor 40U / µL | 80 µL |
| 5 | 0.5mM Spermine | 100mM PMSF | 40 µL |
| 6 | 0.5mM Spermidine | 100mM | 40 µL |
| 7 | 1mM PMSF | 100mM PMSF | 80 µL |
| 8 | 1% Plant Protease Inhibitors | Plant Protease Inhibitors | 80 µL |
| **B** | **1 x GRA\*(total 35 mL)** |  |  |
| 1 | 1× GRAM | 1× GRAM | 35 mL |
| 2 | 1 mM DTT | 1 M DTT | 35 µL |
| 3 | RNase inhibitor 0.4U / µL | RNase inhibitor 40U / µL | 350 µL |
| **C** | **100% Percoll\*(total 5 mL)** |  |  |
| 1 | 100% PM | 100% PM | 5 mL |
| 2 | 1 mM DTT | 1 M DTT | 5 µL |
| 3 | RNase inhibitor 0.4U / µL | RNase inhibitor 40U / µL | 50 µL |
| **D** | **90% Percoll\* (total 5 mL)** |  |  |
| 1 | 90% PM | 90% PM | 5 mL |
| 2 | 1 mM DTT | 1 M DTT | 5 μL |
| 3 | RNase inhibitor 0.4U / μL | RNase inhibitor 40U / μL | 50 μL |
| **E** | **80% Percoll\* (total 5 mL)** |  |  |
| 1 | 80% PM | 80% PM | 5 mL |
| 2 | 1 mM DTT | 1 M DTT | 5 μL |
| 3 | RNase inhibitor 0.4U / μL | RNase inhibitor 40U / μL | 50 μL |
| **F** | **75% Percoll\* (total 5 mL)** |  |  |
| 1 | 75% PM | 75% PM | 5 mL |
| 2 | 1 mM DTT* | 1 M DTT | 5 μL |
| 3 | RNase inhibitor 0.4U / μL | RNase inhibitor 40U / μL | 50 μL |
| **G** | **70% Percoll\* (total 5 mL)** |  |  |
| 1 | 70% PM | 70% PM | 5 mL |
| 2 | 1 mM DTT | 1 M DTT | 5 μL |
| 3 | RNase inhibitor 0.4U / μL | RNase inhibitor 40U / μL | 50 μL |
| **H** | **50% Percoll\* (total 5 mL)** |  |  |
| 1 | 50% PM | 50% PM | 5 mL |
| 2 | 1 mM DTT | 1 M DTT | 5 μL |
| 3 | RNase inhibitor 0.4U / μL | RNase inhibitor 40U / μL | 50 μL |
| **I** | **25% PM\* (total 5 mL)** |  |  |
| 1 | 25% PM | 25% PM | 5 mL |
| 2 | 1 mM DTT | 1 M DTT | 5 μL |
| 3 | RNase inhibitor 0.4U / μL | RNase inhibitor 40U / μL | 50 μL |
| **J** | **1x PBS\* (total 500 μL)** |  |  |
| 1 | 1× PBS (pH=7.4) | 1× PBS (pH=7.4) | 500 μL |
| 2 | 1 mM DTT | 1 M DTT | 0.5 μL |
| 3 | RNase inhibitor 0.4U / μL | RNase inhibitor 40U / μL | 5 μL |
| 4 | 0.04% BSA | 10 mg/ml BSA (1%) | 20 μL |

Reagents marked with \* should be freshly prepared before use.

### Other supplies and Equipment (see Note 7)

1. 15, 50 mL Corning tubes (Thermo-scientific)
2. 2ml low-retention microcentrifuge tubes (Thermo-scientific)
3. 10, 20 μm (PluriSelect), 30 μm (Sysmex) cell strainer
4. Double-edged razor blades
5. Regular 10 μL, 200 μL, 1mL pipette (Eppendorf)
6. Regular 10 μL, 200 μL, 1mL pipette tips (Thermo-scientific)
7. Refrigerated Centrifuge
8. 0.22 μm filter membrance
9. Syringe filter
10. Disposable cell counting chambers
11. Fluorescence Microscope

### Nuclei extraction steps for plant tissues (Illustrated with maize, see Notes 8-15, Fig.1, and Supplementary video)

1. In a pre-chilled 50 mm Petri dish, add 2 mL of freshly prepared and ice-cold 1x Extraction Buffer (1x EB). Immediately place the freshly harvested root tissues into the buffer and finely chop it using a sterile double-edged razor blade for no more than 10-15 minutes, or until a uniform slurry is obtained. Following homogenization, allow the slurry to rest on ice for 5 minutes to stabilize the nuclei before proceeding to filtration (see Note 8).
2. Pre-wet both 30 μm and 20 μm cell strainer with ice-cold 1 × EB to minimize sample loss and improve flow. Filter the crude homogenate first through the 30 μm strainer into a 2 mL low-retention microcentrifuge tube to remove large tissue fragments. Subsequently, pass the filtrate through the 20 μm strainer into a fresh 2 mL low-retention tube to further eliminate small debris and enrich intact nuclei. Keep the filtered nuclei suspension on ice for 5 minutes.
3. In a 15 mL Corning centrifuge tube, first add 1 mL of 70% Percoll. Then, carefully insert the pipette tip to the bottom of the tube and slowly layer 1 mL of 80% Percoll beneath the 70% layer. Using the same method, insert the pipette tip to the bottom of the tube again and slowly layer 1 mL of 100% Percoll beneath the 80% layer (Figure. 1C) (see Note 9).
4. Carefully load the filtered nuclei suspension on top of the 70% Percoll layer without disturbing the gradient.
5. Centrifuge at 1800 × g for 20 minutes at 4^°^ C, using acceleration setting 3 and deceleration setting 3 to preserve gradient integrity (see Note 10).
6. After centrifugation, distinct layers will be visible. The nuclei typically accumulate at the interface between the 80% and 100% Percoll layers, appearing as a milky white band.
7. Carefully remove the second Percoll layer and the milky interface (approx. 900-1000 μL) into a new 15 mL tube (see Note 11). Add at least 6x volume of 1× Gradient Buffer. Centrifuge at 4°C, 700g for 12 min (acceleration 3, deceleration 3).
8. Remove the supernatant, leaving ∼10 μL. Add 50 μL of 1× PBS and mix gently. For counting, mix 10 μL nuclei suspension with 8 μL 1× PBS and 2 μL DAPI (5 ng/ μL), incubate for 1 minute, and load 15 μL into the counting chamber, and count using Fluorescence Microscope.

**Fig. 1.**
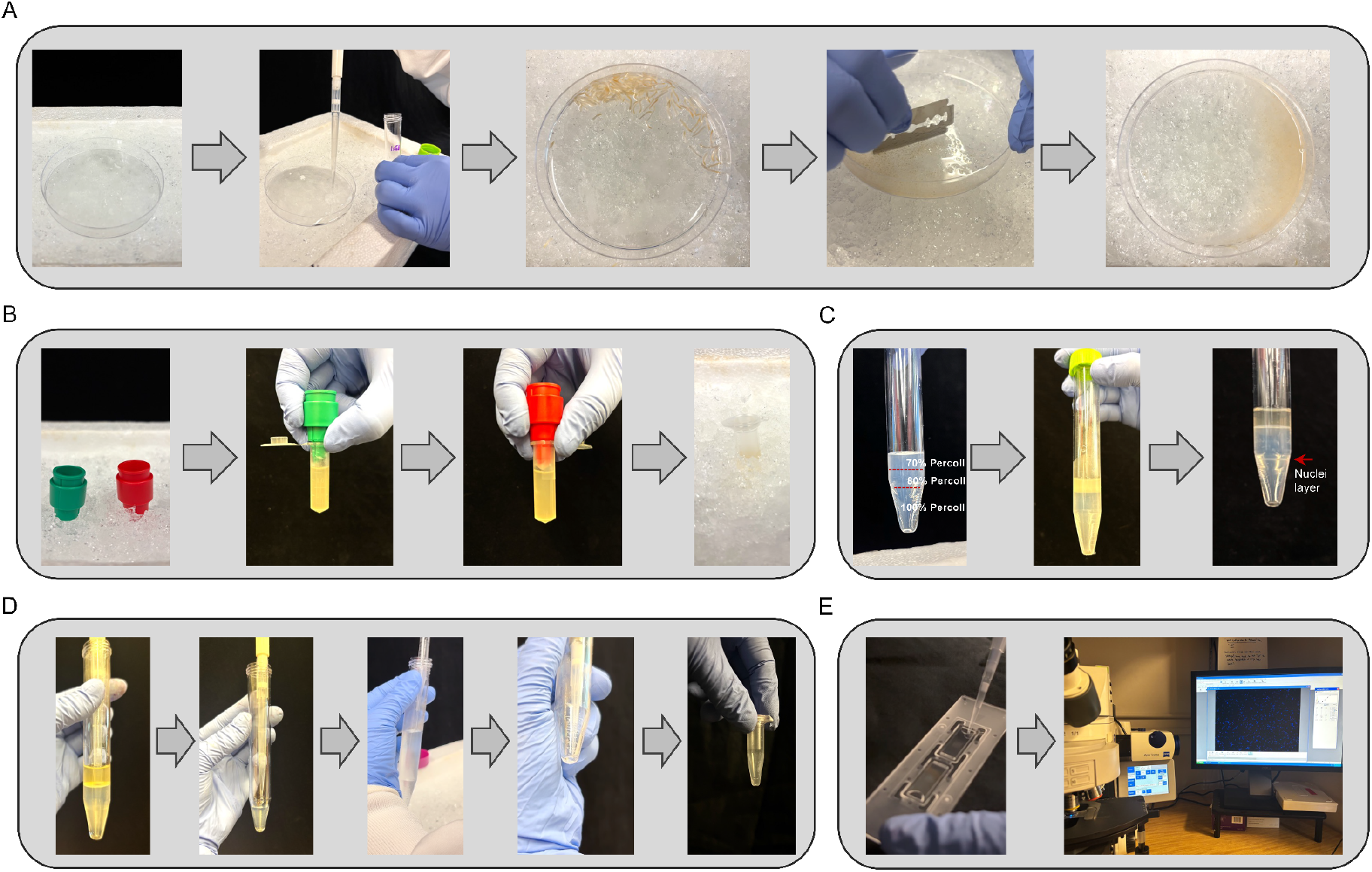
Workflow of nuclei isolation, purification, counting, and quality assessment. (A) Tissue chopping in ice-cold Extraction Buffer (Nuclei extraction step 1). (B) Filtration of the homogenate through 30 μm and 20 μm strainers (Nuclei extraction step 2). (C) Percoll gradient preparation and nuclei enrichment by centrifugation (Nuclei extraction step 3-6). (D) Collection of the nuclei layer, followed by washing and resuspension in ice-cold 1 × PBS solution (Nuclei extraction step 7). (E) Nuclei quality assessment under a fluorescence microscope (Nuclei extraction step 8).

**Notes:**

1. MES buffer (1.25 M, pH 5.6 – 6.7) may crystallize when stored at 4 °C. Before use, warm the solution to 65 °C and mix until fully dissolved to ensure buffer consistency.
2. Spermidine, PMSF, DTT, and DAPI working solutions should be aliquoted and stored at −20 °C, protected from light. Avoid repeated freeze thaw cycles to maintain reagent stability and activity.
3. PMSF (100 mM) should be prepared in anhydrous ethanol, stored at -20 °C in amber tubes, and used within one week. It is highly unstable in aqueous solution – prepare fresh immediately before use whenever possible.
4. Percoll should be stored at 4 °C. Before use, gently invert the bottle several times to ensure even distribution of colloidal silica particles; do not vortex or shake vigorously.
5. Pre-extraction buffer stock solutions (e.g., 4× EBM) are stable at 4 °C for up to one month. However, working solutions (e.g., 1× EB) should be prepared fresh daily to ensure optimal nuclei preservation and compatibility with downstream applications.
6. RNase inhibitor (40 U/μL) is relatively expensive and can be omitted during initial protocol optimization without compromising nuclear integrity. However, for samples intended for sequencing, it is strongly recommended to include RNase inhibitors during nuclei isolation to minimize RNA degradation.
7. All plasticware (tubes, tips, filters) must be RNase-free and sterile to avoid contamination and degradation of nuclear RNA, especially when isolating nuclei for transcriptomic analyses.
8. During chopping, ensure the tissue remains fully submerged in 1× EB to minimize shear stress and oxidative damage. Chopping time should be limited to 10 – 15 minutes to preserve nuclear envelope integrity. Over-chopping may lead to chromatin leakage, nuclear rupture, and increased debris.
9. Percoll gradients are essential for effective nuclei enrichment. After layering 70%, 80%, and 100% Percoll, clearly defined interfaces should be visible. Layer each solution slowly along the tube wall or using a pipette tip positioned at the bottom, avoiding air bubbles or pressure that could disrupt the gradient.
10. Use a swinging-bucket refrigerated centrifuge with slow acceleration and deceleration settings (e.g., speed setting 3/3). Sudden acceleration or mechanical vibration can disrupt the Percoll gradient, reducing nuclei recovery and purity.
11. If isolated nuclei contain visible debris or cell fragments, an additional filtration step using a 10 μm or 20 μm strainer after Step 7 can be performed to improve sample purity prior to downstream applications.
12. It is recommended that all reagents and consumables (e.g., tubes, filters) be pre-chilled and kept on ice during the nuclei isolation process to preserve nuclear integrity.
13. Gloves and masks should be worn at all times to prevent contamination from RNAse, DNAse, and protease activities present on the hands and others.
14. The volumes and the number of tubes used here are ideal for 0.5g-1g of tissue. If more tissue is used, this can create problems in overloading the homogenization and miracloth filtration stages, and overloading on the Percoll gradients. If more nuclei are required, it’s best to carry out several preparations in parallel or use larger apparatus.
15. The optimal conditions for nuclei isolation can vary substantially across species and tissue types, and may require empirical fine-tuning. Based on our optimization experience, we recommend the following: (1) Initial gradient selection can be guided by genome size. A preliminary test using three relatively wide-spaced gradient layers allows estimation of the nuclei’s buoyant position. This can then be refined to determine the optimal gradient composition for the target species or tissue; (2) The final concentration of Triton X-100 in the lysis buffer influences membrane disruption, nuclei release, and nuclear integrity. If lysis is inefficient or nuclei rupture is excessive, adjust the Triton X-100 concentration in the initial extraction buffer; (3) In tissues rich in secondary metabolites, supplementing the extraction buffer with 0.1% PVP can improve nuclei yield and purity; (4) Both speed and acceleration/deceleration rates can impact nuclei recovery and integrity. If abundant intact nuclei are present before enrichment but show damage or reduced yield after Percoll centrifugation, consider optimizing the centrifugation profile (e.g., stepwise acceleration/deceleration) to improve recovery while maintaining integrity.

## Results

### Validation of nuclei isolation from maize multiple tissues

To evaluate the performance of the nuclei isolation protocol across different maize tissues, we collected four types of samples: approximately 60 root tips (1-2 cm) from 4-day-old germinated seeding, 0.5 g of stem tissue from the same seedings, 0.5 g of leaf tissue from 12-day-old seedings, and approximately 30 embryos harvested 20 days after pollination. All samples were subjected to nuclei isolation using protocol described above, followed by Percoll density gradient centrifugation. For maize samples, a discontinuous Percoll gradient consisting 70 %, 80 %, and 100 % layers was used. Nuclei enrichment at the 80 % Percoll layer was consistently observed across all tissue types, appearing as a distinct milky-white band, which indicated successful separation and high nuclei purity (Fig. 1).

Then to assess nuclei integrity and purity, isolated nuclei were stained with DAPI and examined by fluorescence microscopy (Fig. 2). In the brightfield images, nuclei were observed as intact, round to oval structures. Under the DAPI channel, these nuclei showed strong blue fluorescence, indicating preserved nuclei DNA. A small proportion of non-nuclear particles were also visible under brightfield and showed weak DAPI staining, likely due to nonspecific DNA binding. By integrating information from both brightfield and fluorescence images, we reliably distinguished intact nuclei from debris and quantify nuclear yield. For maize root tips, stem, leaf, and embryo samples, the nuclei concentrations were 1440, 1280, 1400 and 1360 nuclei μL^**-1**^, respectively. Given the 40 μL suspension volume per sample, each preparation contained > 50,000 nuclei, meeting the input requirements for downstream single-nucleus RNA-seq.

**Fig. 2.**
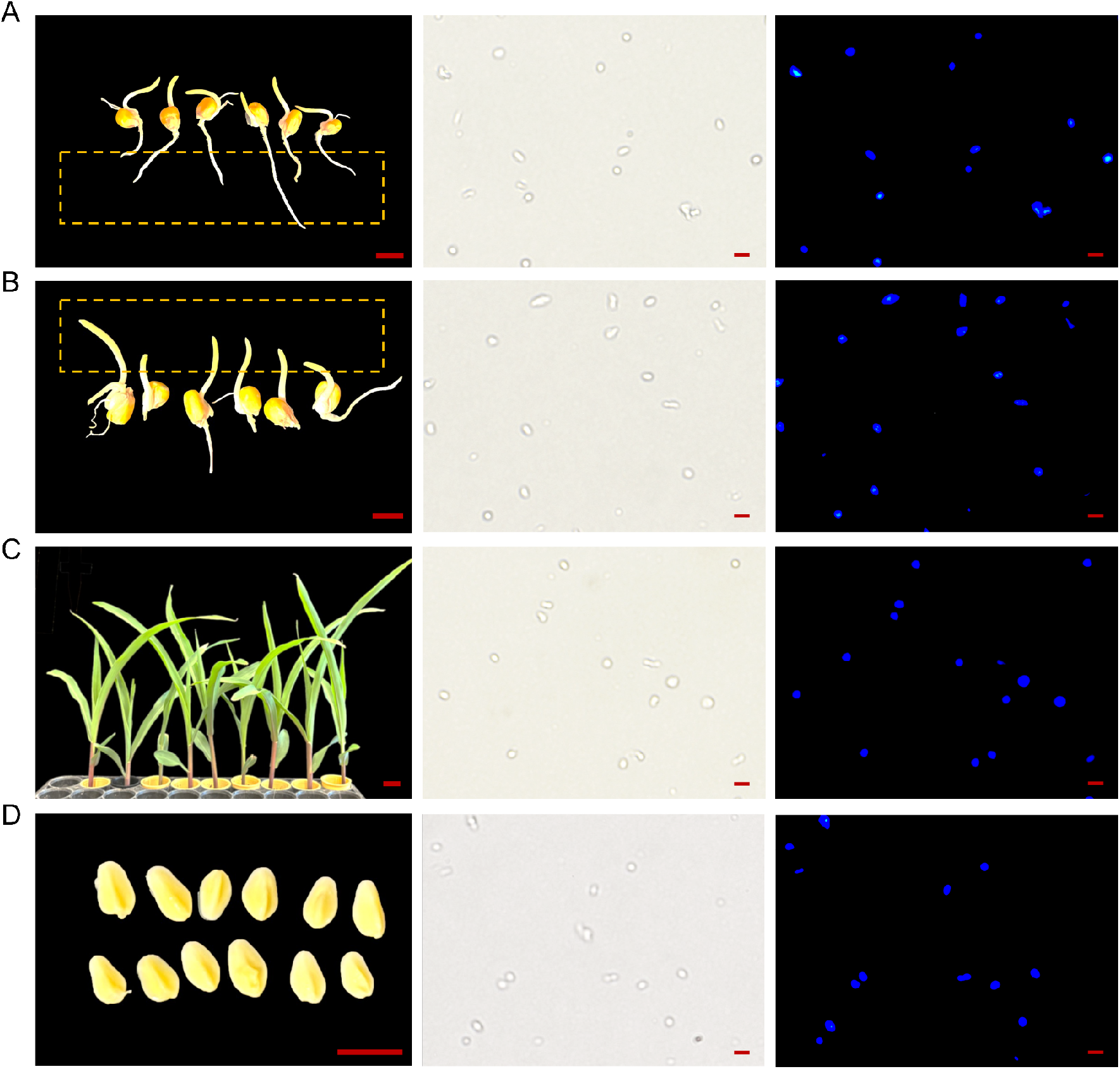
Nuclei morphology visualized by brightfield and DAPI fluorescence microscopy in maize shoot (A), root (B), leaf (C), and embryo (D). The middle column shows brightfield images of isolated nuclei, while the right column shows corresponding DAPI-stained fluorescence images. Scale bars: 1 cm (left column); 10 μm (middle and right columns).

### Optimization of nuclei isolation across plant species

We successfully isolated nuclei from root tips and leaf tissues of A*rabidopsis*, rice, sorghum, blue grama, *Tripsacum*, barley, oat, and wheat using the optimized protocol. Intact nuclei were consistently recovered across all species and tissue types. To accommodate the wide range of genome sizes, we adjusted the Percoll gradient composition accordingly (Fig. 3). For *Arabidopsis* (genome size ∼0.135 Gb), a two-layer gradient (25% and 75%) was used, with nuclei enriched in the 25% layer. For species with moderate genome sizes, including rice, sorghum, and blue grama (∼0.43–1.08 Gb), a three-layer gradient (25%, 50%, and 75%) was applied, and nuclei accumulated at the 50% layer. For *Tripsacum* (∼5.1 Gb), a higher-density three-layer gradient (70%, 90%, and 100%) was used, with nuclei enriched in the 90% layer. In contrast, for large-genome species such as barley, oat, and wheat, a single-layer 25% Percoll gradient was applied, and nuclei were recovered at the bottom of the tube.

**Fig. 3.**
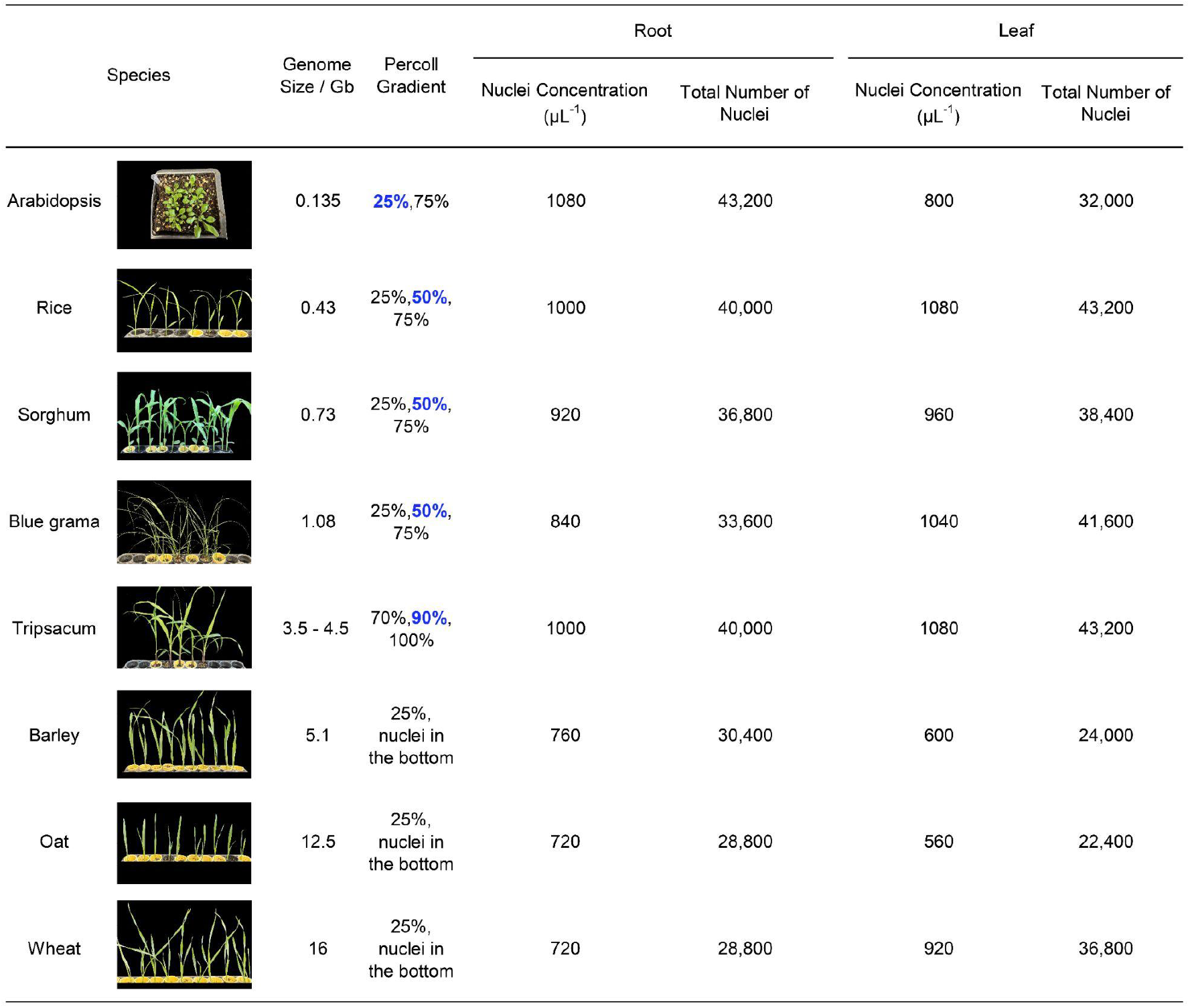
Comparison of nuclei isolation across 8 plant species with varying genome sizes and optimal Percoll gradients. The Percoll gradient used for each species is listed, along with nuclei concentration and total number. Blue text indicates the Percoll layer nuclei predominantly enriched.

Nuclei isolated from Arabidopsis, rice, sorghum, blue grama, and *Tripsacum* using the multilayer gradients showed high purity with minimal background debris, and yields typically ranged from 22,400 to 43,200 nuclei per sample (Fig. 3, Supplementary Fig.1 and 2). Although similar yields were obtained from barley, oat, and wheat using the single-layer method, the resulting suspensions contained more background debris. For these species, we recommend consulting the sequencing provider regarding the need for further purification. If necessary, sucrose gradient centrifugation or fluorescence-activated nuclei sorting (FANS) can be employed to improve sample quality for downstream applications [33].

### Assessment of nuclei integrity and cDNA quality for downstream snRNA-seq

To further validate our protocol, we selected leaf-derived nuclei from species of particular interest - maize and *Tripsacum* – and submitted them to 10x Genomics Chromium platform to assess nuclei quality and downstream compatibility. Nuclei integrity and concentration were first evaluated with the Invitrogen Countess II Automated Cell Counter. In this assay, GFP fluorescence marks residual intact cells or lipid-rich debris, while TxRed fluorescence specifically labels nuclear DNA. Both maize and *Tripsacum* suspensions contained a high proportion of TxRed^+^ intact, uniformly sized nuclei with minimal debris or aggregates. Samples were diluted threefold prior to counting, yielding measured concentrations of 1248 nuclei μL^−1^ for maize and 1146 nuclei μL^−1^ for *Tripsacum* (Fig. 4A, B), corresponding to over 40,000 nuclei per sample, sufficient for high-throughput library preparation.

**Fig. 4.**
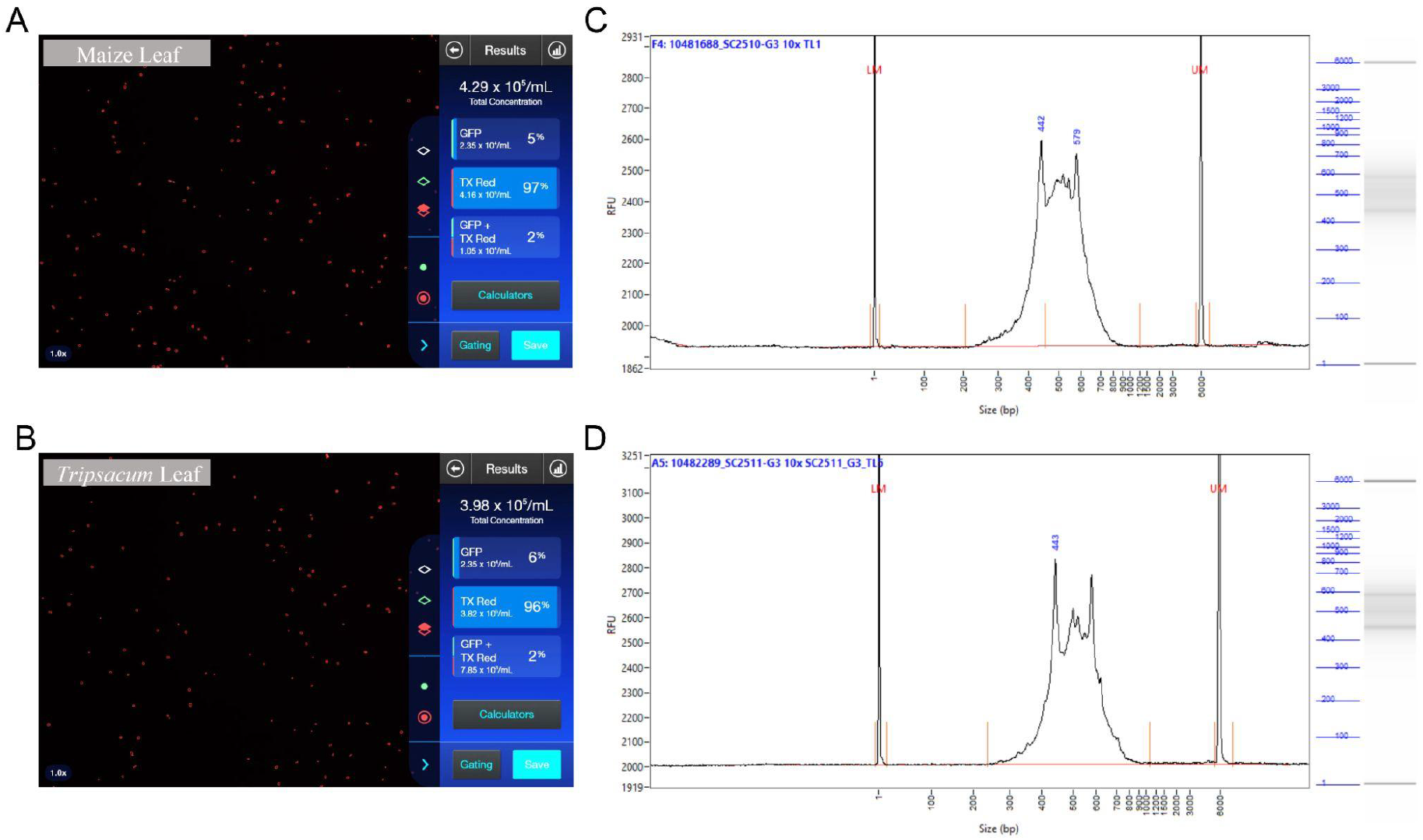
Nuclei concentration and cDNA quality assessment for maize and *Tripsacum* leaf samples. (A, B) Nuclei concentration and quality measured using Countess 3 Automated Cell Counter for maize (A) and *Tripsacum* (B). (C, D) Bioanalyzer electropherogram traces showing cDNA library quality from maize (C) and *Tripsacum* (D) nuclei samples. The graphs display library size distribution after cDNA amplification.

Following nuclei encapsulation and reverse transcription, cDNA quality was assessed using an Agilent Bioanalyzer. Both species displayed sharp, distinct cDNA size distribution peaks corresponding to full-length transcripts, with minimal low-molecular-weight smear, indicating high RNA integrity and efficient reverse transcription (Fig. 4C, D). These results confirm that nuclei prepared with our protocol are of high quality and fully compatible with robust snRNA-seq library construction, supporting its broad applicability across species.

## Conclusions

Together, our results demonstrate that the nuclei isolation protocol we developed is broadly effective across diverse plant species and tissue types. By tailoring Percoll gradient compositions according to genome size, we consistently obtained high-quality nuclei from roots, leaves, stems, and embryos across nine species, including both monocots and dicots. The protocol enabled the recovery of intact nuclei with minimal debris and sufficient yield (20,000–50,000 nuclei per sample), as confirmed by DAPI staining and fluorescence imaging. Furthermore, nuclei derived from maize and *Tripsacum* leaf tissues were successfully used for single-nucleus RNA-seq library preparation on the 10x Genomics platform, yielding sharp cDNA profiles with minimal degradation. These findings validate the robustness, flexibility, and downstream compatibility of the protocol, providing a reliable workflow for plant single-nucleus transcriptomic studies across species with diverse genome sizes and cellular complexity.

## Acknowledgements

This work was supported by a cooperative agreement from the U.S. Department of Agriculture Agricultural Research Service to Cornell University (MCR) and by the National Natural Science Foundation of China (Grant No. 32301840). We thank Zongyan Liu (Cornell University) for assistance with video recording and Charlie Hale (Cornell University) for providing the background music.

## Author contributions

Yun Luo designed this study. Jiali Yan developed the standard nuclei extraction protocol and Yun Luo optimized the protocol. Yun Luo and Thuy La performed the nuclei extraction experiments in different species. Yun Luo wrote the draft and revised the manuscript. All authors read and approved the manuscript.

## Data availability

No datasets were generated or analyzed during this study.

---

**Supplementary Fig. 1.**
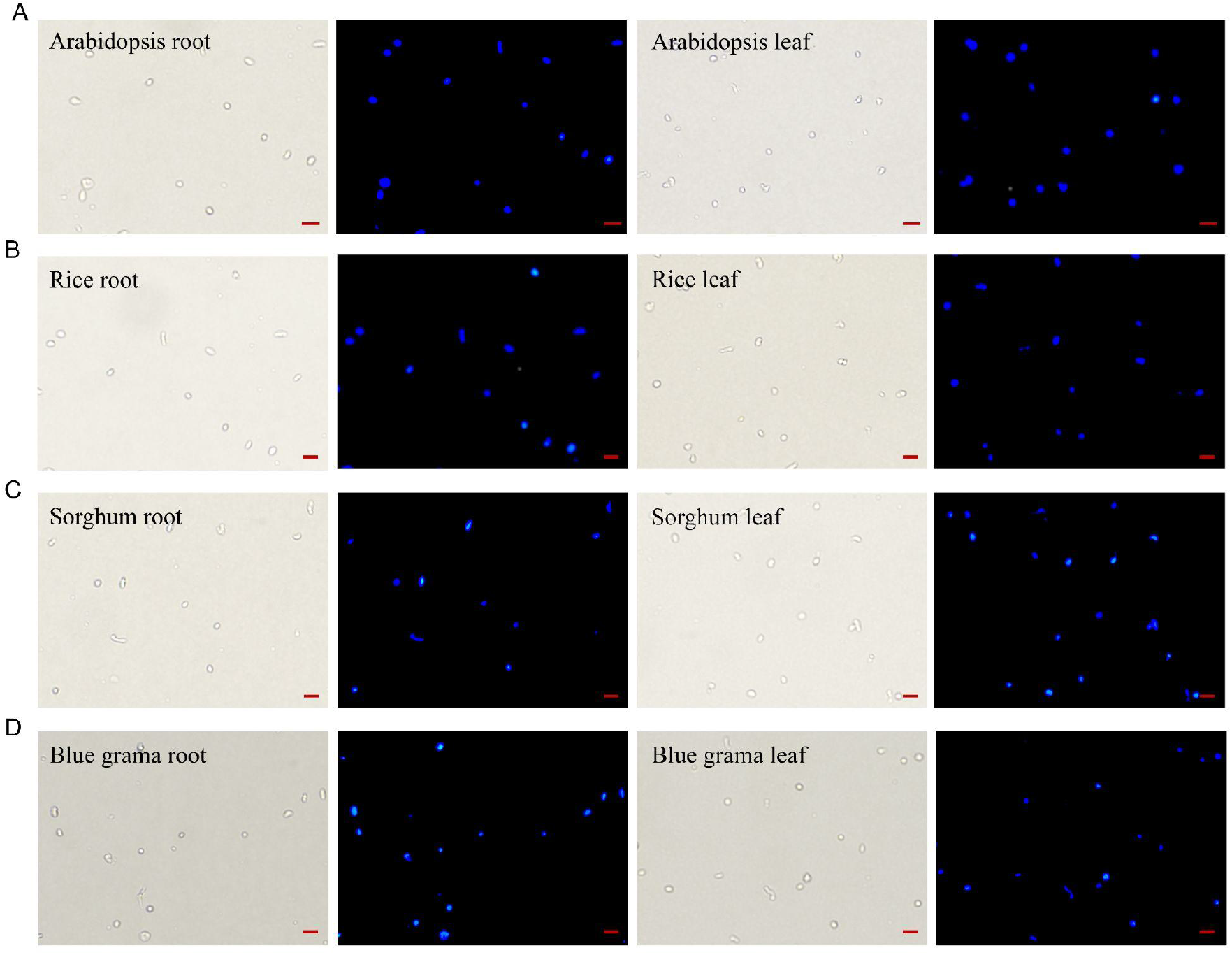
Representative images of isolated nuclei under brightfield and DAPI fluorescence microscopy in Arabidopsis, rice, sorghum and blue grama root and leaf. Scale bar = 10 μm.

**Supplementary Fig. 2.**
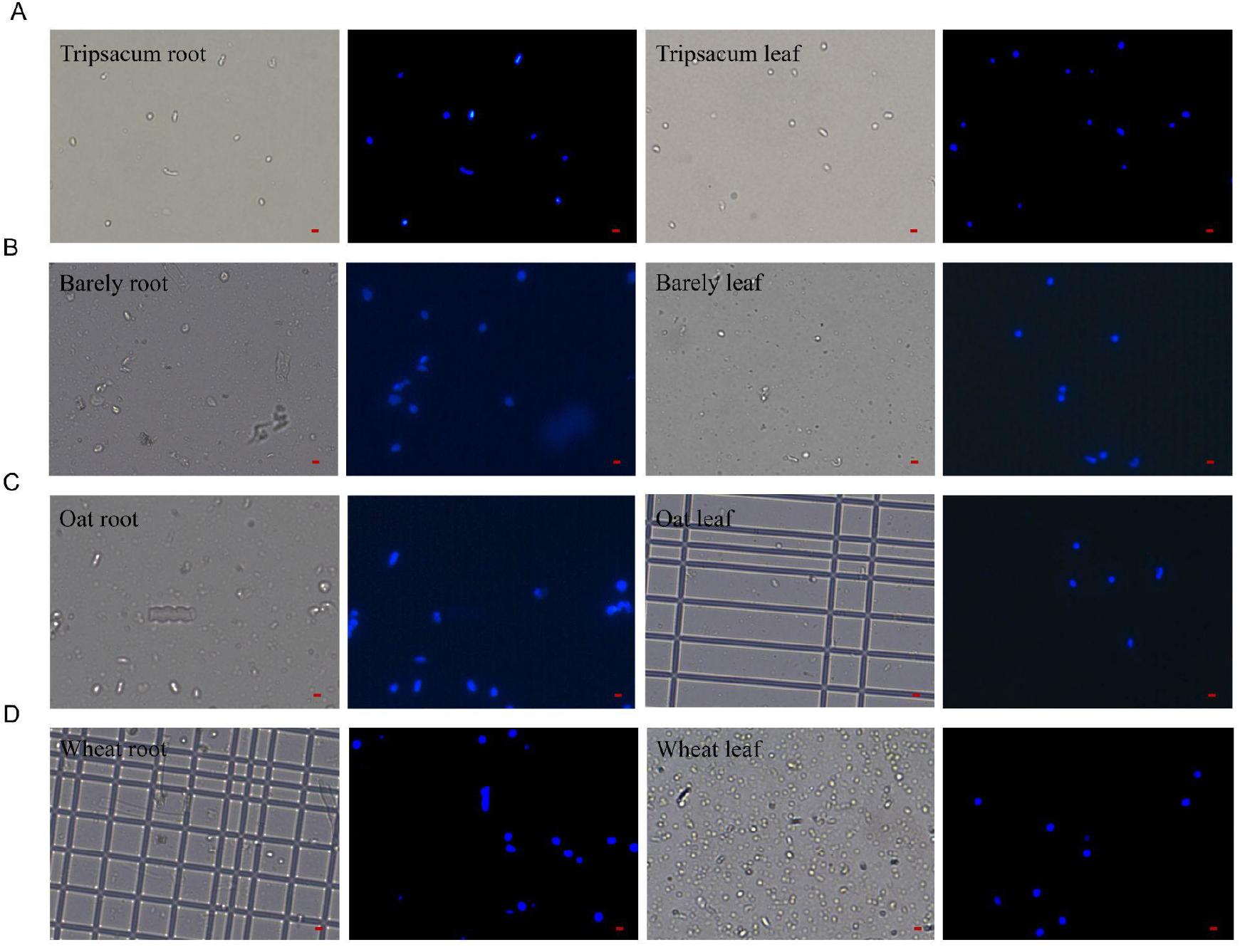
Representative images of isolated nuclei under brightfield and DAPI fluorescence microscopy in Tripsacum, barely, oat and wheat root and leaf. Scale bar = 10 μm.

